# Discrete Distributional Differential Expression (D^3^E) - A Tool for Gene Expression Analysis of Single-cell RNA-seq Data

**DOI:** 10.1101/020735

**Authors:** Mihails Delmans, Martin Hemberg

## Abstract

The advent of high throughput RNA-seq at the single-cell level has opened up new opportunities to elucidate the heterogeneity of gene expression. One of the most widespread applications of RNA-seq is to identify genes which are differentially expressed between two experimental conditions. Here, we present a discrete, distributional method for differential gene expression (D^3^E), a novel algorithm specifically designed for single-cell RNA-seq data. We use synthetic data to evaluate D^3^E, demonstrating that it can detect changes in expression, even when the mean level remains unchanged. Since D^3^E is based on an analytically tractable stochastic model, it provides additional biological insights by quantifying biologically meaningful properties, such as the average burst size and frequency. We use D^3^E to investigate experimental data, and with the help of the underlying model, we directly test hypotheses about the driving mechanism behind changes in gene expression.

## Background

Over the last two decades, several methods for global quantitative profiling of gene expression have been developed [24, 30, 39]. One of the most common uses of gene expression data is to identify differentially-expressed (DE) genes between two groups of replicates collected from distinct experimental conditions, e.g. stimulated vs unstimulated, mutant vs wild-type, or at separate time-points. The goal of DE analysis is to identify genes that underlie the phenotypical differences between the conditions.

The first method for genome-wide expression profiling was microarrays, but as sequencing costs have decreased, profiling by direct sequencing of the transcriptome (RNA-seq) has become more popular. Initially, RNA-seq experiments were carried out in bulk on samples of up to 10^5^ cells. Consequently, only information about the mean expression of each gene in a sample could be extracted. However, it has been known since the 1950s [23] that gene expression varies from cell to cell, and more recently it has been shown that stochastic variation may play an important role in development, signaling and stress response [28, 29, 41]. Thus, recently developed single-cell RNA-seq protocols [16, 37], could potentially provide a greater understanding of how the transcriptome varies between cells with the same genotype and cell-type. The main advantage of single-cell RNA-seq over bulk RNA-seq is the fact that one obtains the full distribution of expression levels, rather than the population mean. To take full advantage of single-cell data, for DE analysis as well as for other types of investigation, e.g. inference of gene regulatory networks, novel analysis methods are required.

Single-cell DE analysis is complicated by the fact that comparison of two probability distributions is an ambiguous task. With the exception of SCDE [18], most common tools for preforming single-cell DE analysis - DESeq2 [21], Cuffdiff [38], limma [32] and EdgeR [33] - are all adaptations of bulk RNA-sequencing methods. They mainly focus on filtration and normalisation of the raw data, and DE genes are identified based on changes in mean expression levels. The main drawback of using only the mean is that one ignores the gene expression heterogeneity, and will thus fail to detect situations where, for example, there is only a change in the variance of gene expression. Alternative methods for comparing probability distributions are the Kolmogorov-Smirnov test, the likelihood ratio test and the Cramér-von Mises test. What these methods have in common is that they summarize the difference between two distributions as a single value, which can be used to test for significance.

Gene expression at the single-cell level has been studied theoretically for almost three decades [4], and the most widely used model is referred to as the transcriptional bursting model. The transcriptional bursting model [25, 27] provides a mechanistic description of the stochastic switching of the promoter as well as the the production and degradation of transcripts at the single cell level (Fig. 1A,B). The model makes it easy to generate synthetic data which closely resembles the experimental data. Since the model is analytically tractable it allows us to derive several other biologically relevant properties of gene expression (Fig. 1C) [19], which makes it easier to interpret results biologically. Despite its simplicity, the transcriptional bursting model enjoys strong experimental support [8, 19, 36, 42, 43].

**Figure 1:**
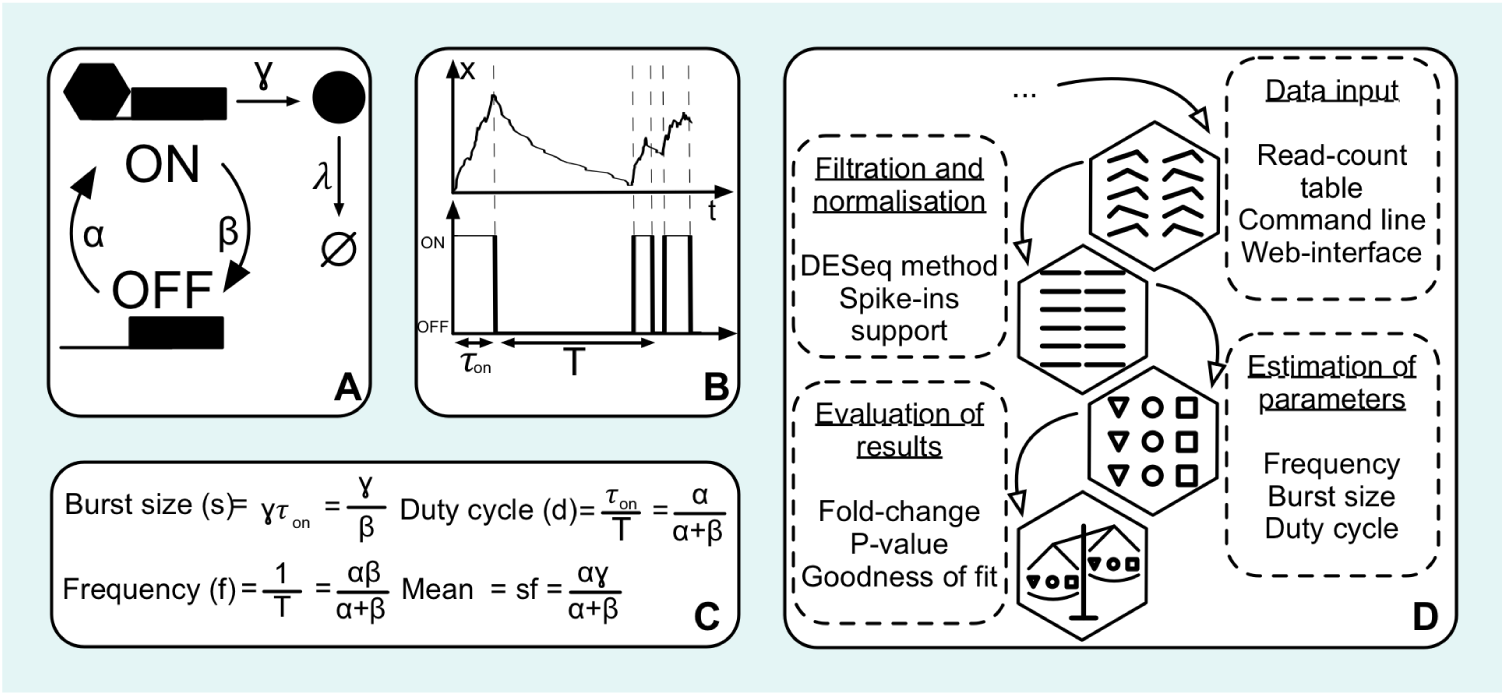
Overview of D^3^E. **A)** Graphical representation of the transcriptional bursting model. **B)** Example of a realization of the transcriptional bursting model with parameters *α* = 1, *β* = 10, *γ* = 100, and *λ* = 1 [12]. In this regime, the gene exhibits a bursty behavior with a bimodal stationary distribution. **C)** Derivation of the biologically-relevant parameters from the parameters of the transcriptional bursting model. **D)** Flowchart of the D^3^E algorithm.

In this paper, we present D^3^E, a method based on the comparison of two probability distributions for performing differential gene expression analysis. D^3^E consists of two separate modules: a module for comparing expression profiles using the Cramér-von Mises, the likelihood-ratio test, or the Kolmogorov-Smirnov test, and a module for fitting the transcriptional bursting model. Thus, D^3^E allows the user to go beyond merely identifying DE genes and provides biological insight into the mechanisms underlying the change in expression. We demonstrate the power of D^3^E to detect changes in gene expression using synthetic data. Finally, we apply D^3^E to experimental data to demonstrate its ability to detect significant changes which are not reflected by the mean.

## Results and Discussion

### Algorithm and Implementation

D^3^E takes a read-count table as an input, with rows and columns corresponding to transcripts and cells, respectively. The user should split the columns into two or more groups by providing cell labels in the input file. If there are more than two groups of cells, they must be compared one pair at a time. D^3^E uses four steps to process the data. First, input data is normalised using the same algorithm as DESeq2 (see *Methods*) and filtered by removing the genes that are not expressed in any of the cells. Second, the Cramér-von Mises test, the Kolmogorov-Smirnov (KS) test, or the likelihood ratio test [11] is used to identify the genes with a significant change in expression between the two samples of interest. Third, the transcriptional bursting model is fitted to the expression data for each gene in both samples using either the method of moments or a Bayesian method [19]. Fourth, the change in parameters between the two samples is calculated for each gene (Fig. 1D).

A command-line version of D^3^E written in Python can be downloaded from GitHub (https://hemberg-lab.github.io/D3E), and the source code is available under the GPL licence. Furthermore, there is also a web-version available at http://www.sanger.ac.uk/sanger/GeneRegulation_D3E. Due to the time required to run D^3^E, the web version limits the number of genes and cells that may be analyzed, and it can only use the method of moments for estimating parameters.

### DE Analysis Module

To compare distributions obtained from two different sets of cells, D^3^E uses either the Cramér-von Mises test, the KS test or the likelihood ratio test to quantify the difference in gene expression (see *Methods*). The first two tests are non-parametric which is advantageous since it allows us to apply D^3^E to any single-cell dataset, not just the ones collected using RNA-seq. The null hypothesis for all three tests is that the two samples are drawn from the same distribution. The premise of D^3^E is that when two samples are drawn from the same population of cells, the test should result in a high *p*-value. On the other hand, if the cells are drawn from two populations with different transcript distributions, then the resulting *p*-value should be low.

We first evaluated D^3^E using synthetic data. Fortunately, there is a widely used, experimentally validated stochastic model available for singlecell gene expression [25]. We refer to this model as the transcriptional bursting model (Fig. 1A), and it is characterized by three parameters: *α*, the rate of promoter activation; *β*, the rate of promoter inactivation; *γ*, the rate of transcription when the promoter is in the active state; and a transcript degradation rate *λ*. The stationary distribution of the transcriptional bursting model takes the form of a Poisson-Beta mixture distribution [19, 25] 

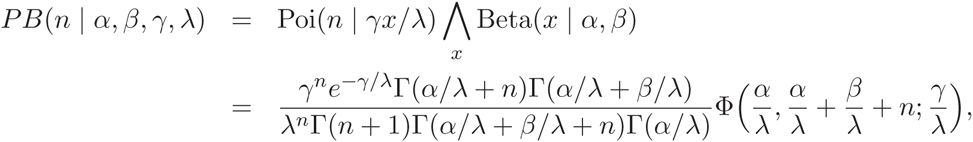
where *n* is the number of transcripts of a particular gene, *x* is an auxiliary variable, Γ is the Euler Gamma function, and Φ(*a*, *b*; *c*) is the confluent hypergeometric function.

An important feature of the Poisson-Beta distribution is that the three parameters *α*, *β* and *γ* are normalised by the rate of mRNA degradation *λ*. Consequently, when fitting the parameters for the Poisson-Beta distribution from a stationary sample, only three parameters can be estimated, and they are unique up to a common multiplicative factor, *λ*. Since a single-cell RNA-seq experiment corresponds to a snapshot of individual cells, it is often reasonable to assume that the samples are drawn from the stationary distribution. The inability to uniquely identify all four parameters from single-cell RNA-seq data means that it is only approprite to apply the transcriptional bursting model to DE analysis when *λ*s are constant between the compared samples, or when the degradation rates are known for both samples. Without knowledge of *λ* it is impossible to unambiguously determine how the parameters have changed.

To evaluate the sensitivity of the Cramér-von Mises, the KS and the likelihood ratio test to changes in the parameters, we selected triplets of parameters (*α*, *β*, *γ*) from a range that is characteristic for single-cell RNA-seq data [16]. For each parameter triplet one of the parameters was varied, while fixing the remaining two, and a series of tests was carried out on the corresponding Poisson-Beta samples. For each combination of parameters, we assumed that there were 50 cells from each condition when generating the data. The results can be summarized by a set of matrices, where rows and columns correspond to values of the varied parameter, and the elements in the matrix are *p*-values from the tests (Fig. 2C). Ideally we would like to find high *p*-values on the diagonal and low *p*-values away from the diagonal. We used a heuristic for characterizing the pattern of *p*-values, and for each matrix we obtained a single score, *S* (see *Methods*). When *S* = 0, high *p*-values are only found on the diagonal, suggesting that D^3^E has successfully identified genes where there was a change in one of the parameters.

**Figure 2:**
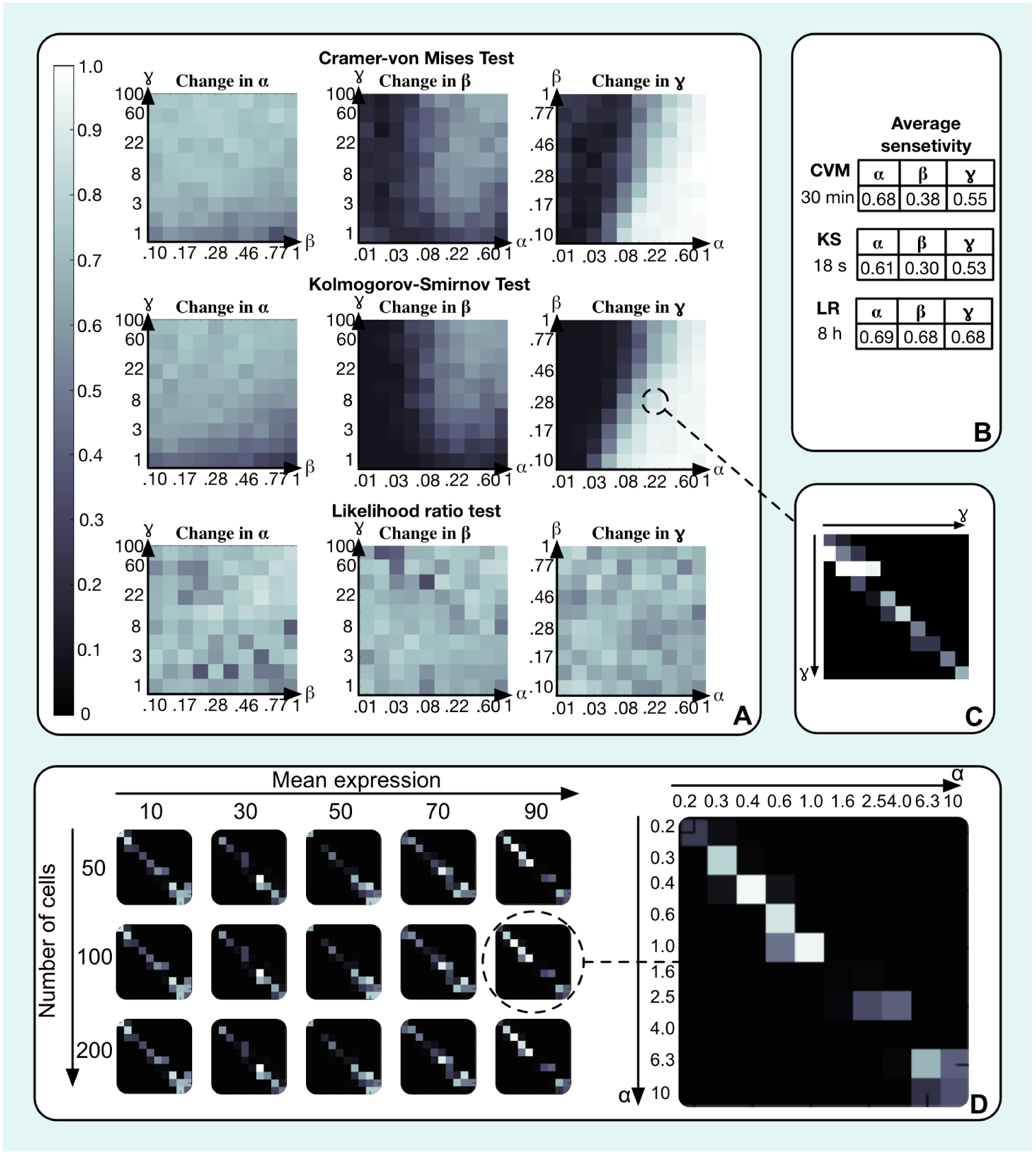
DE analysis for synthetic data. **A)** Sensitivity to changes in parameters of the Poisson-Beta distribution for the Cramér-von Mises, KS and likelihood ratio tests. A lighter color denotes a low sensitivity to changes of a particular parameter. **B)** Average sensititivity scores for the panels in A as well as the run-times. **C)** An example of a matrix which was used to assign the colors in **A**. Here, parameters *α* = .22 and *β* = .28, while *γ* is varied from 1 to 100 on a log-scale. Each element in the matrix reflects a *p*-value of a KS test between two Poisson-Beta distributions with the corresponding parameters. We expect to find high values along the diagonal, where the changes are smaller. **D)** DE analysis for the scenario where the mean is fixed but the variance is changed. D^3^E is able to reliably identify differentially expressed gene based on the change in the shape of distribution alone. Here, the color represents the *p*-value for the Cramér-von Mises test with dark colors indicating a low value.

The results suggest that all three tests are capable of accurately detecting changes in the parameters in certain regions of the parameter space (Fig. 2A). For all three tests, changes in *β* are the most difficult to detect while changes in *γ* are the easiest to identify. The methods perform poorly when *γ* is small and either *β* is large or *α* is small. In this regime, the Poisson-Beta distribution is similar to the Poisson distribution with a mean close to zero, and it is challenging to identify which parameter has changed, and by how much. From a biological perspective, when a transcription rate is small and a gene has a small duty cycle (small *α* or big *β*) there are almost no transcripts produced since the promoter spends most of its time in the inactive state. Therefore, changes in either of the three parameters will be difficult to distinguish. The performance of each method can be summarized by the average score across all combination of parameters and we find that the likelihood ratio test is the most accurate, followed by the Cramér-von Mises test and the KS test (2B, S1). The accuracy, however, is also mirrored by the computational costs; analyzing the data in Fig. 2A takes about 18 seconds for the KS-test, 30 minutes for the Cramér-von Mises test and 8.3 hours for the likelihood ratio test using a MacBook Pro laptop with a 3.2 GHz Intel i5 processor and 16 Gb of RAM.

We also considered the scenario when the two distributions are different, but the mean is identical. This is a situation where it is all but impossible for methods which only use the mean to reliably detect that there has been a change in the expression profile. In contrast, D^3^E is able to reliably identify a change in expression. Our results show that a change of *α* and *β* by a factor of 2, which is roughly equivalent to changing the variance by the same factor, is sufficient for the *p*-value to drop below .05 for a sample of 50 cells (Fig. 2D).

A particular challenge for DE analysis is to determine the *p*-value threshold for when a change can be considered significant. The traditional approach is to use a fixed value, e.g. .05, and then adjust for multiple hypothesis testing. For D^3^E two different methods are offered for selecting the critical *p*-value. On the one hand, the user can specify a false discovery rate, and the tool will use the Benjamini-Hochberg procedure [6] to identify the critical *p*-value. Alternatively, one may take an empirical approach whereby one of the two samples is first split into two parts. By definition, the two parts should have identical distributions for all genes, which means that it can be used as a negative control. D^3^E applies one of the three tests to the negative control, and records the lowest *p*-value identified, *p*^*^. When comparing the two original distributions, only genes with a *p*-value below *a* × *p*^*^ are considered significant, where the default value for the parameter *a* is .1. We advice the user that the reduced sample size of the negative control set is likely to result in a less stringent cut-off than what would be expected if the negative control had the same sample size as the original data. To evaluate the heuristic strategy, we generated 1,000 pair of samples with the same number of reads and cells, using identical parameter values for the samples in each pair. Using the Cramér-von Mises test we recorded the lowest observed *p*-value, *p*^*^, and we used .1 × *p*^*^ as a threshold for when to call a test significant. For both the method of moments and the Bayesian method, we found that 97% of the genes were detected as not DE. The control experiment demonstrates that D^3^E is capable of accurately distinguishing situations where the parameters are unchanged.

To further evaluate the performance of D^3^E relative to other DE methods, we generated additional synthetic data sets where one of the three parameters was varied while the other two were fixed as before. For each data set we designated genes as DE where the parameter had changed by at least a factor of 1.5, 2 or 4. The arbitrary decision of what constitutes a significant change allows us to define the calls of the DE algorithms as either true positive, false positive, true negative or false negative. The results can be summarized as a ROC curve, and again we find that changes in *β* are more difficult to detect compared to the other two parameters (Fig. 3). Importantly, we find that for larger parameter changes, D^3^E is always amongst the best performing methods (Fig. 3).

**Figure 3:**
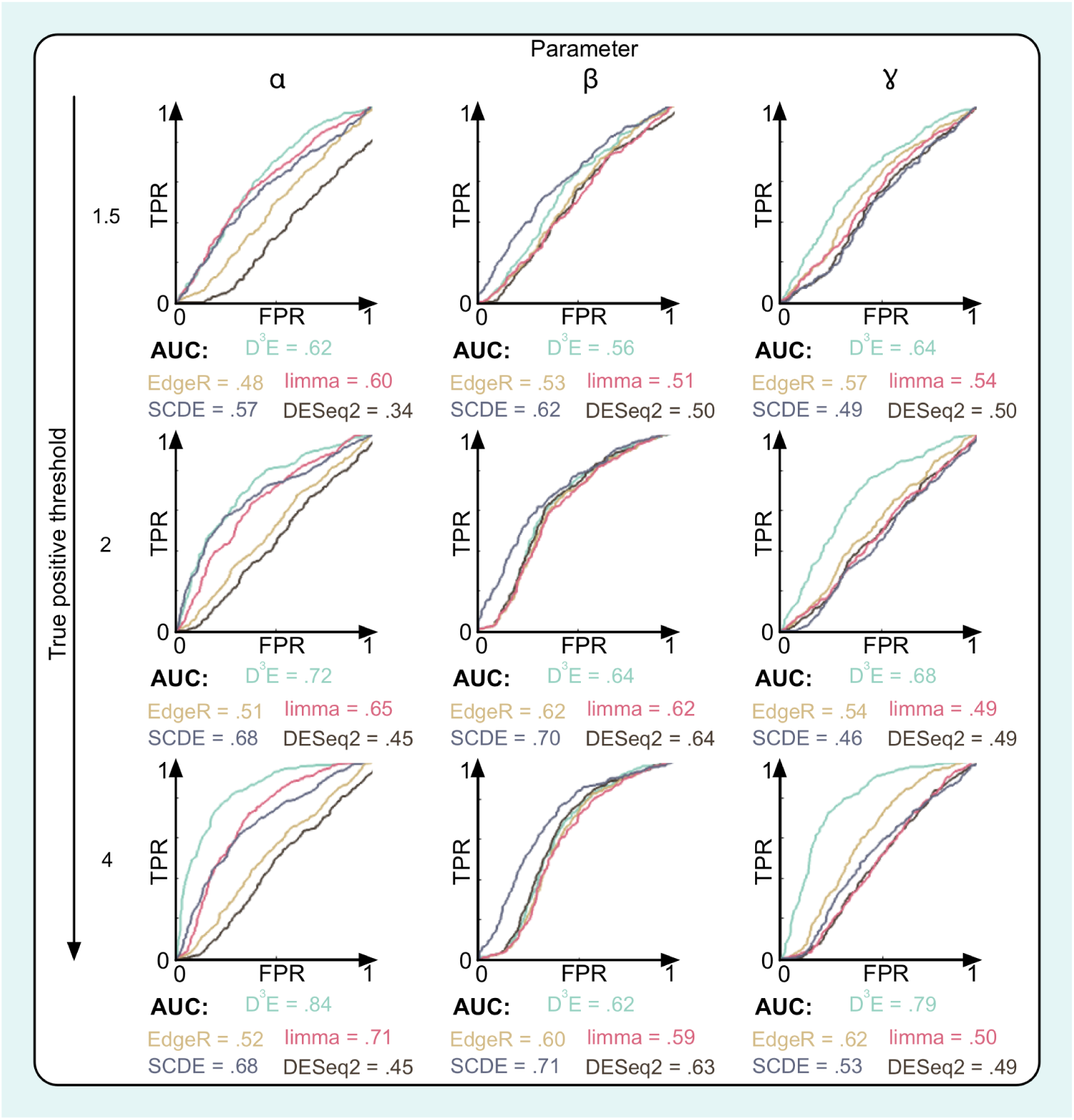
Comparison of DE methods for synthetic data. Each panel shows the receiver operator characteristics (ROC) calculated for synthetic data using five different DE algorithms. The numbers below each panel indicate the area under the curve. The rows correspond to different thresholds for when a gene is considered significantly changed. DESeq2 reports `NA` for many genes. Since the NA cannot be interpreted as either DE or not DE, we treat these calls as false, which explains the unusual shape of the ROC curve and the fact that the AUC value is below .5.

### Parameter Estimation Module

The likelihood ratio test is a parametric test, and thus it requires estimates of the parameters *α*, *β* and *γ*. D^3^E has two approaches; the method of moments which is relatively fast and a more computationally costly Bayesian inference method (see *Methods*) [19]. We evaluated the accuracy of the parameter estimates, both assuming perfect sampling as well as in the scenario when some transcripts are lost due to low starting levels of mRNA [18]. In our simulations, we first generated parameters for the Poisson-Beta model by drawing from a distribution similar to the one for the data reported by Islam *et al* [16]. We then assumed that the dropout probability scaled with the mean expression level as 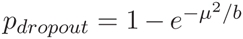 [26], where the parameter b controls the rate of dropouts. In our simulations we used *b* = 10, 100 and 200. Our simulations show that the relative error for the estimates of the parameters *α*, *β* and *γ* are relatively robust to dropout events (Fig. 4A).

**Figure 4:**
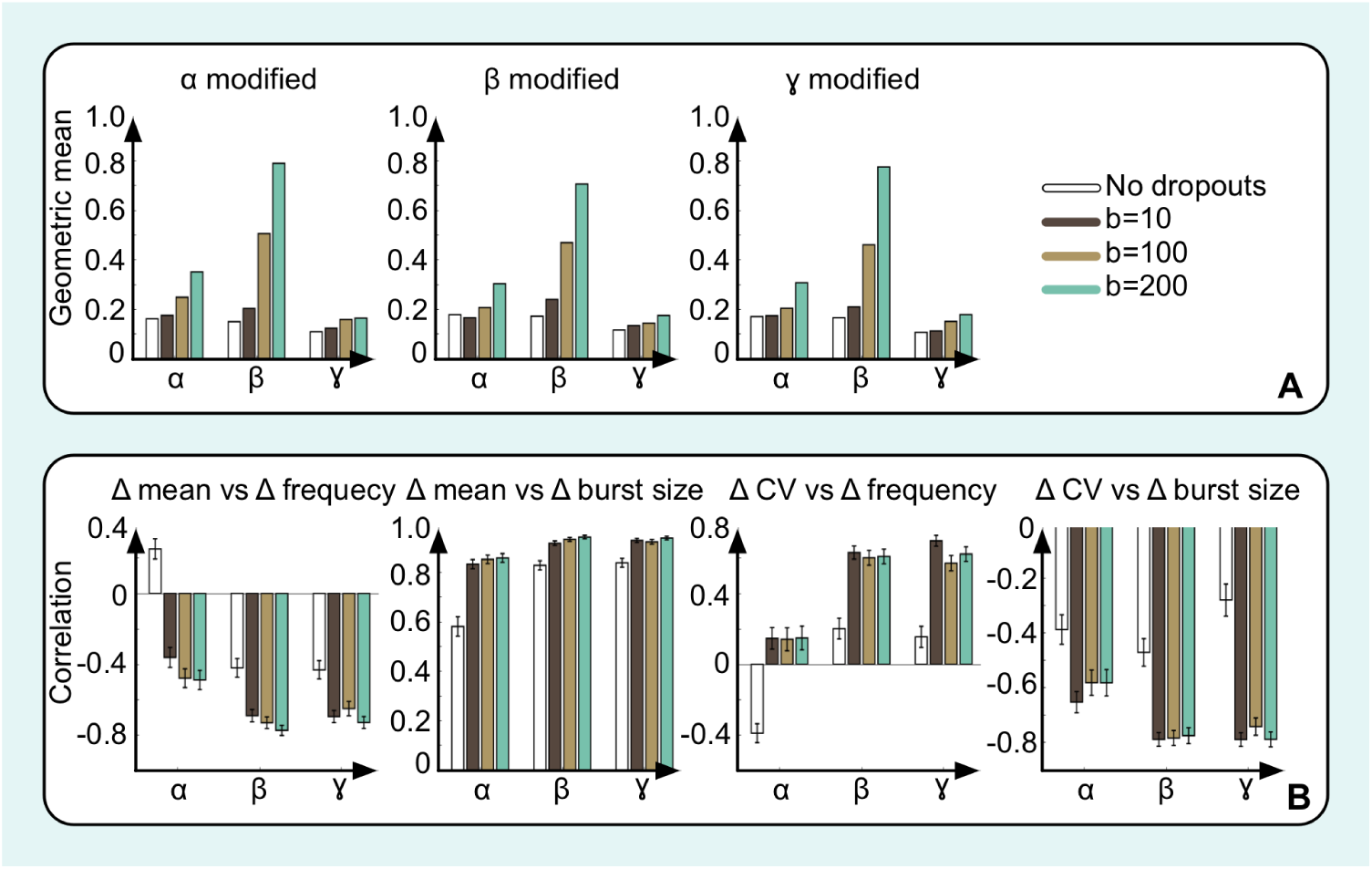
Sensitivity to transcript dropout errors. **A)** Errors for the estimates of the parameters for the synthetic data in Fig. 2A using the Bayesian inference method. Each bar represents the geometric mean squared relative error for the parameter estimates **B)** Errors for the estimates of the correlations between the estimated changes in parameters for the synthetic data in Fig. 2A. Each bar represents the estimated Pearson correlation coefficient between the log-changes of the quantities.

An important advantage of using the transcriptional bursting model (Fig. 1) is that it is possible to derive other quantities - the average burst size, the burst frequency, the mean expression level, and the proportion of time in the active state (duty cycle) - which are easier to measure and interpret biologically than the parameters *α*, *β* and *γ*. Importantly, the transcriptional bursting model allows us to learn more about *how* the expression level has changed between the two conditions. In the transcriptional bursting model, there are three different ways to increase the mean expression level; by decreasing the degradation rate, by increasing the burst frequency, or by increasing the burst size. By comparing the correlations between the change in mean expression levels and the change in burst size or burst frequency, it is possible to gain additional biological insights relating to what aspect has led to the change in gene expression. In contrast to the estimates of the rate parameters the correlation estimates are sensitive to dropout events (Fig. 4B), and one must thus be careful when interpreting the results.

### Application to Experimental Data

The tests on synthetic data suggest that D^3^E can reliably identify differentially expressed genes. A more useful test of the algorithm, however, involves experimental data which has been reliably validated. Unlike bulk data [30], unfortunately there are no gold-standard datasets available. Nonetheless, to further evaluate D^3^E, we considered the single-cell RNA-seq data from two and four-cell mouse embryos where qPCR data from the same cell-types was collected for 90 genes [5]. Unfortunately, the correlation of changes in gene expression between the qPCR and RNA-seq data (*ρ*_Δ_ = .46) (Fig. S2) is even worse than the correlation of the individual samples (*ρ*_2_ = .6, *ρ*_4_ = .5). Thus, it does not come as a surprise that the overlap between the genes which are considered DE in the qPCR experiment has little overlap with genes which are considred DE from RNA-seq by any of the five algorithms that we compared (Table S1). Even so, we find large differences in the number of genes identified as DE, ranging from 1 (edgeR) to 35 (DESeq2).

To further evaluate D^3^E, we applied it to the two datasets collected by Islam *et al.* [16] from 48 mouse embryonic stem cells and 44 mouse embryonic fibroblasts. To establish the *p*-value threshold, we first separated the stem cells into two groups, and compared the expression (see *Methods*). We used this approach for determining the threshold for D^3^E, SCDE, edgeR and limma, while for DESeq2, we used the adjusted *p*-value reported by the software. When comparing the two cell-types, D^3^E identified 4716 genes as DE, DESeq2 identified 6360 genes, limma-voom identified 7245 genes, edgeR identified 1140 genes, and SCDE identified 1092 genes (Fig. 5D). Surprisingly, the agreement between the five methods is quite low with only a core set of 380 genes identified by all three methods. If we require a gene to be identified of 4 out of 5 methods, then an additional 495 genes are idenitified as DE, suggesting that there is a set of around 900 genes which can confidently considered DE. To further evaluate the set of genes identified as DE by each method, we investigated the distiribution of fold-change values (Fig. 5A). The distributions gives an indication of how large fold changes are required for detection, and we note most of the genes have a higher expression in fibroblasts compared to stem cells. Compared to DESeq2, SCDE and edgeR, we also notice that D^3^E is able to identify genes with a lower fold change. Indeed, there were several examples of genes where the change in mean expression was modest, but they were still identified by D^3^E as differentially expressed (Fig. 5C).

**Figure 5:**
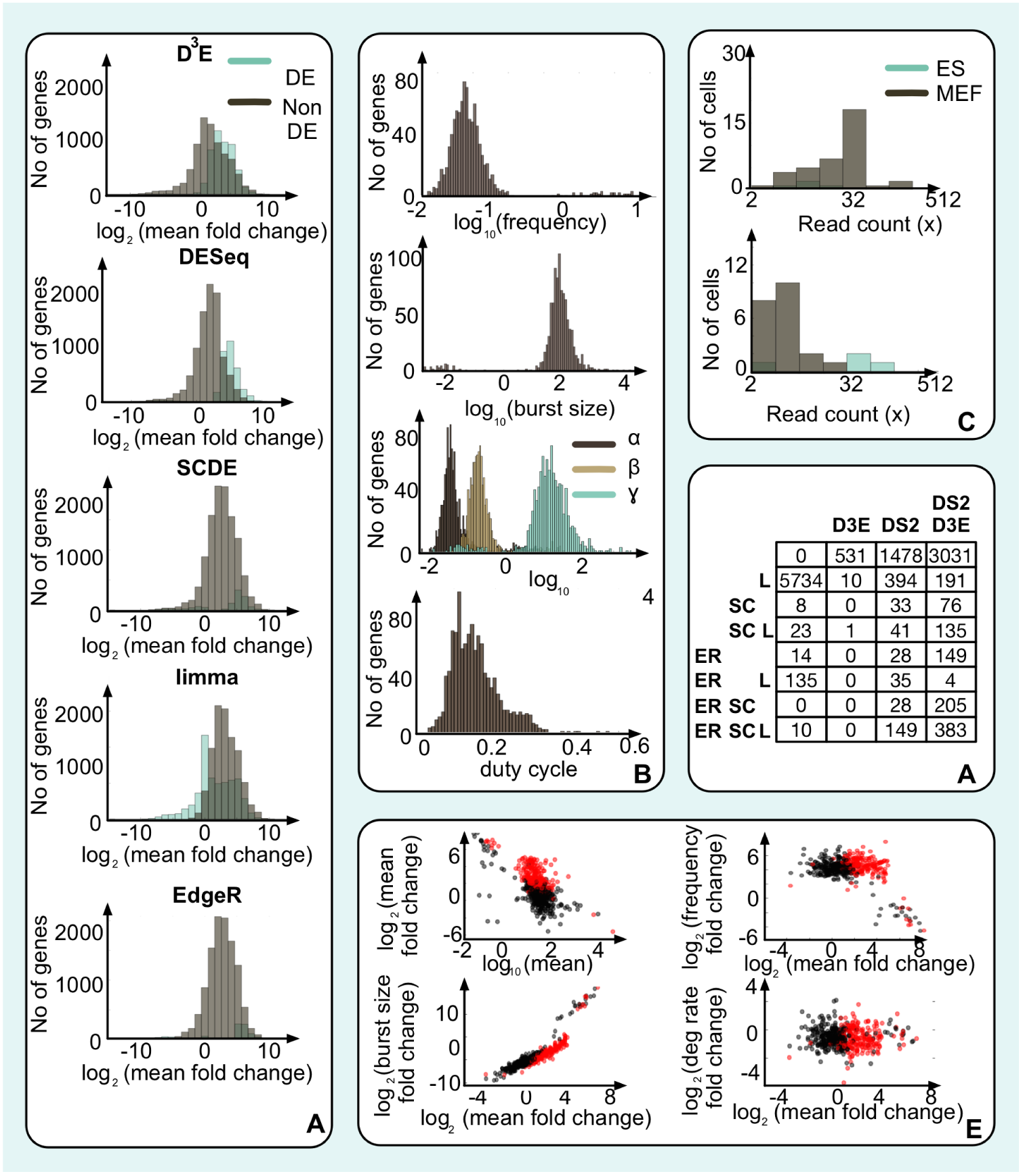
Analysis of experimental data. **A)** Histogram showing the fold-changes for the genes which were considered significantly changed (blue) and not changed (gray) for D^3^E, DESeq2, limma, edgeR and SCDE. **B)** Histograms showing the distribution of parameter values for all cells from [16]. From top to bottom, the panels represent the frequency, the burst size, the inferred rates for the transcriptional bursting model, and the duty cycle. **C)** Examples of two genes, Cdc42bpb in the top panel and Hist1h2bb in the bottom panel, which were identified as DE by D^3^E. In both cases, the change in mean expression is less than 70% whereas the variance increases by > 10-fold. **D)** Karnaugh table showing the number of genes identified as differentially expressed by D^3^E, SCDE, limma, edgeR, and DESeq2 for the two datasets collected by Islam *et al* [16]. **E)** Scatterplots showing the mean in mESCs, and the fold-change, as well as the fold-change of the mean compared to the change in degradation rate, burst frequency and burst size. In all panels, black dots represent genes which did not change, red dots represent genes which were deemed significant by D^3^E.

Next, we took advantage of the transcriptional bursting model underlying D^3^E, and we fitted the parameters *α*, *β*, and *γ* for all genes. We found that for 85% of the genes, at least one of the parameters changed by at least 2-fold, suggesting that there are substantial differences between the two cell-types. The results show that all three parameters follow log-normal distributions, spanning approximately one or two orders of magnitude in both cell-types (Fig. 5B). With the exception of the duty cycle which is constrained to be in the interval [0, 1], the derived quantities showed a similar distribution.

We calculated the three derived quantities for each condition for the 2105 genes where we were able to obtain degradation rates for both cell-types [34, 35]. Next, we compared the changes in degradation rate, burst frequency and burst size to the change in mean expression level (Fig. 5E). The results clearly demonstrate that it is the change in burst size which underlies the change in mean expression levels (*ρ* = .91), suggesting that altering the burst size is the driving mechanism behind differences in mean expression between conditions. However, considering our simulation experiments (Fig. 4B), it is likely that the true correlation is lower.

Another property of interest is the coefficient of variation (CV), defined as the standard deviation divided by the mean, which is used to quantify the gene expression noise. The CV is inversely correlated with the mean, and the transcriptional bursting model reveals that the change in CV is mainly correlated with the change in the duty cycle (*ρ* = .47), while the effect of a change in burst size is considerably smaller (*ρ* = .24, Fig. S3). To further demonstrate the use of the transcriptional bursting module, we also investigated changes in the auto-correlation times of each gene. The auto-correlation provides information about the time-scale of the noise, i.e. how quickly the gene expression level varies. The expected value of the autocorrelation, *τ_c_*, is given by (*Methods*)

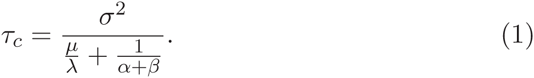

Comparison of *τ_c_* and the change in the mean for the Islam *et al* data reveals that the two quantities are strongly correlated (*ρ* = .87, Fig. S4). However, when investigating all the quantities on the right hand side of Eq. (1) the comparison shows that it is the change in variance which is most strongly correlated with the change in autocorrelation times (*ρ* = .90, Fig. S4).

We also applied the methodology to cells from the early and late blastocyst from mouse embryos [10]. Since we do not have access to degradation rates for these cell-types, not all of the correlations can be explored. Nevertheless, the correlations that can be calculated are largely similar to the ones observed for the Islam *et al* data (Fig S5–6). Taken together, these results demonstrate that it is possible to generate testable hypotheses about how changes in the property of a gene has come about.

### Discussion

DE analysis is one of the most common uses of bulk RNA-seq, and we expect that it will become an important application for single-cell RNA-seq as well. Here, we have presented D^3^E, a tool for analyszing DE for singlecell data. The main difference between D^3^E and other methods is that D^3^E compares the full distribution of each gene rather than just the first moment. Therefore, it becomes possible to identify genes where the higher moments have changed, with the mean remaining constant. To the best of our knowledge, D^3^E is the first method for DE analysis which considers properties other than the first and second moment of the distribution. Using synthetic data, we demonstrate that D^3^E can reliably detect when only the shape, but not the mean is changed.

One of the main challenges in developing a DE analysis method for single-cell RNA-seq data is that, unlike for bulk data, there are no gold-standards available [30]. Comparison of qPCR and RNA-seq data revealed only a modest correlation between the two methods, implying that the two methods are inconsistent. Thus, one must resort to synthetic data for evaluation. Fortunately, for single-cell gene expression, there is an analytically tractable transcriptional bursting model available which has been experimentally validated. Even with synthetic data, however, it is not obvious how one should define a change in expression. Consider the situation where one of the parameters changes by a small amount which is just sufficient to be detected given the limits of the technical noise, the read depth and the sample size. Then the question is whether or not the change is sufficient to be biologically meaningful. Another challenge stems from the difficulty of disentangling the technical and the biological noise. The transcriptional bursting model does not account for the technical noise in single-cell experiments which can be considerable [3, 7, 13, 31]. Since each cell can only be sequenced once, one cannot carry out technical replicates in the same way that can be done for bulk experiments.

Given the lack of ground truth and the difficulties involved in rigorously defining change in gene expression at the single-cell level, we recommend that more than one algorithm is used to identify DE genes. If the aim is to identify the genes which are most likely to have changed significantly, we believe that a consensus approach is the best one to use. Such a strategy will minimize the number of false positives with the drawback of increasing the number of false negatives.

D^3^E implements two different non-parametric methods and one parametric method for comparing probability distributions. The methods emphasize different aspects of the distributions, and they are also associated with different computational costs. An important future research question is to determine what method is the most appropriate for single-cell DE analysis. Since the results are sensitive to technical noise, such developments should ideally be carried out taking the specific details of the protocol into consideration.

We have shown that the transcriptional bursting model makes it possible to extract additional, biologically relevant results from the DE analysis. However, to be able to fully utilize the transcriptional bursting model, the mRNA degradation rates must be known, or assumed to be constant. Determining degradation rates directly remains experimentally challenging, and today they are only available for a handful of cell-types. However, alternative strategies have been proposed, whereby degradation rates are estimated from RNA-seq data using distribution of reads along the length of a gene [14, 40]. The RNA-seq based methods make it possible to estimate degradation rates without further experiments, and they could thus significantly expand the range of samples where the transcriptional bursting model can be applied. Another restriction of our method is that the groups of cells must be known in advance. In many cases, e.g. when comparing samples from a mutant and wild-type or when comparing different stages of development [5], it is straightforward to assign labels to cells. However, there are also situations, e.g. when analyzing a tissue-sample, when the cell-labels are unknown and in these scenarios D^3^E is no longer applicable.

## Conclusions

Our work combines three important aspects of genomics - high-throughput sequencing technologies, computational data analysis, and systems biology modelling. In the present study, we have combined single cell RNA-seq, non-parametric comparison of distributions and an analytical model of stochastic gene expression which allows us to extract biologically meaningful quantities, providing insights not just about which genes have changed between two conditions, but also how the change has come about.

## Materials and Methods

### Cramér-von Mises criterion

To compare two empirical distributions of read counts from different cell samples, the Cramér-von Mises test was used. Let *x*_1_, *x*_2_,... *x_N_* and *y*_1_, *y*_2_, . . . *y_M_* be the observed read counts for the two samples. Given ranks *q_i_* and *s_i_* of the read-counts from the first and the second samples, in the ordered pooled sample, the Cramér-von Mises criterion is given by [1]:

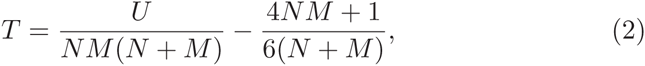

where

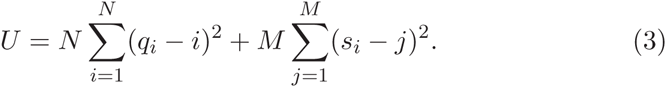

The p-value associated with a null-hypothesis that two samples are drawn from the same distribution was calculated as [2]:

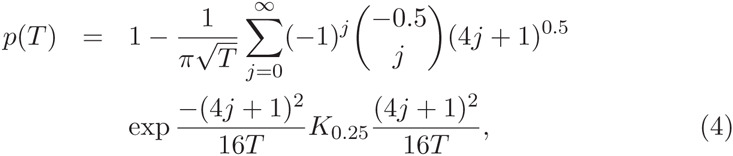

where

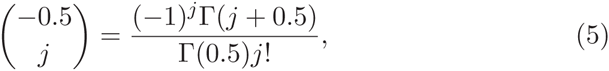

Γ(*z*) is Euler’s Gamma function, and *K_v_*(*z*) is a modified Bessel function of the second kind.

The infinite sum in (4) converges fast after the first few terms. In practice, the *p*-value was calculated using first 100 terms of the sum for values of *T* less or equal to 12. For values of *T* greater than, 12 the *p*-value was set to zero.

### Parameter estimation

A fast but inaccurate method for estimating parameters of a Poisson-Beta distribution is a moments matching technique. The parameters can be expressed through the sample exponential moments [25]:

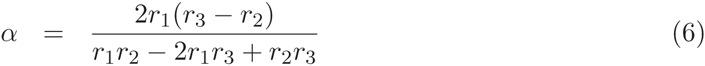

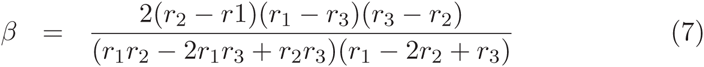

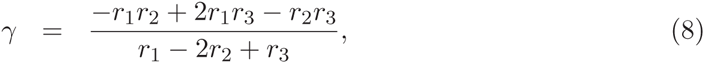
where *r_i_* is a successive ratio of exponential moments *e_i_*: 

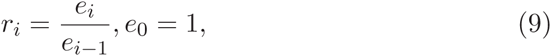
for an i’th exponential moment: *e_i_* = E[*X*(*X* – 1)...(*X* – *i* + 1)], where *X* is a sample of read counts.

The parameters of a Poisson-Beta distribution can also be estimated by a Bayesian inference method [19]. The Bayesian method is more accurate, but it requires more computational power. A Gamma distribution was used as a prior for the parameters *α*, *β* and *γ*:

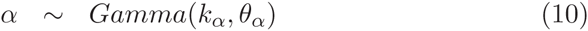

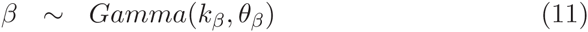

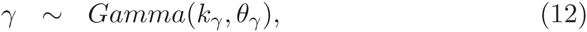
where 

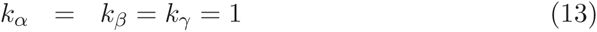

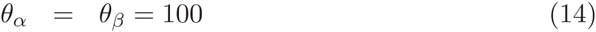

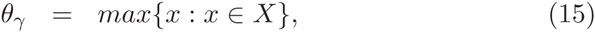

The number of read counts, x, was drawn from a Poisson-Beta distribution:

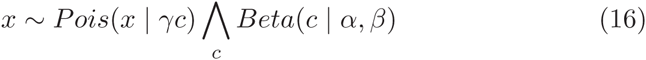

Parameter estimation was performed by a collapsed Gibbs sampler, using Slice sampling [22]. Conditional distributions for parameters during sampling were given by:

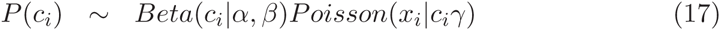

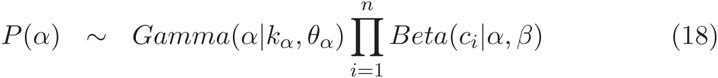

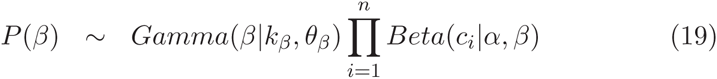

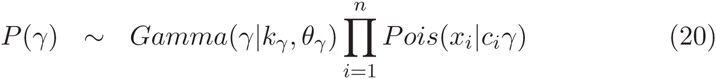

### Likelihood ratio test

The likelihood ratio test provides a parametric test for differential expression. One of the two conditions is designated as the control and the other is designated as the test conditions. For each gene, the log-likelihood of the data from the test condition is calculated using the parameters estimated for both the control and the test. We then test the null hypothesis that the ratio of the likelihoods calculated using the two parameters is not significantly different from 1.

Calculating the likelihood using the Poisson-Beta distribution numerically is very challenging due to the presence of the confluent hypergeometric function [19]. The methods available in Python become unreliable for large values of the third argument. To be able to evaluate the Poisson-Beta distribution we employ a Monte Carlo method whereby the PDF is estimated using *N* randomly generated samples. *N* can be set by the user as a flag and the default value is 1,000.

### Synthetic data

Synthetic data was produced by sampling from a Poisson-Beta distribution, i.e. first drawing an auxiliary variable *c* from Beta distribution with parameters *α* and *β*: *c* ~ *Beta*(*α*, *β*) and then drawing from a Poisson distribution with parameter *λ* = *cγ*: *x* ~ *Poisson*(*cγ*).

### Analyis of the sensitivity

To evaluate how well D^3^E is capable of detecting changes in different regimes of the parameter space, we systematically varied the three parameters of the Poisson-Beta model across the range of values representative of the biological data, *α* ∈ [.01, .1], *β* ∈ [.1,1], and *γ* ∈ [1,100]. We fixed a pair of parameters and varied the third in 10 steps over its range, recording the *p*-value for one of the tests. For each of the 100 different combinations, it was assumed that the sample consisted of 50 cells from each condition was generated. Close to the diagonal, the changes in the parameters are small, and we expect a high *p*-value in these positions. To summarize the matrix of *p*-values, we calculate a composite score as 

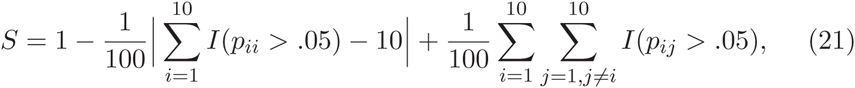
where *I* is the indicator function. The score S ranges between 0 and 1, and 1 is obtained when all of the *p*-values on the diagonal are greater than .05 while the off-diagonal elements are smaller than .05. The score is mapped to a color and reported in Fig. 2.

### Dropout analysis

To evaluate the effect of experimental noise, we considered the possibility of transcript dropout events. Dropouts are likely to occur either as a consequence of failure to isolate the transcript when lyzing the cell, failure of the RT reaction, or failure of the amplification. The probability of failure is poorly understood, but it has been suggested that the dropout probability can be modeled as 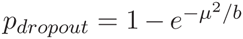, where *μ* is the mean expression and *b* is a parameter [26].

For our dropout experiments, we randomly generated 1,000 parameter triplets based on the fitted data from Islam *et al* [16]. For each triplet, we changed one parameter at a time and the change was allowed to be up to 4-fold. Next, we generated realizations from both distributions for 50 cells in each condition. We used the parameter estimation module to evaluate the accuracy of the estimates for perfect samples as well as for three different values of the dropout parameter *b* = 10, 100, 200.

### Normalization

The normalization of the raw read counts was performed by the same method used by DESeq2 [21]. Let *x_ij_* represent the raw number of reads for *i* = 1,2..*N* and *j* = 1,2..*M*, where *N* is the number of genes, and *M* is the total number of cells in the experiment. Then, the size factor *s_j_* is found as

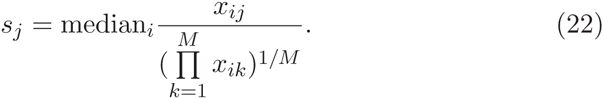

The corrected read counts are then calculated as 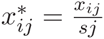. The size factors are calculated based on spike-ins data only if it is available.

### Determining *p*-value threshold

To determine the *p*-value threshold for D^3^E, we first take the sample which will be used as the control group (i.e. in the denominator when calculating the fold-change), and split it into two non-overlapping subsets. Next, one of the tests is applied to the split sample, and the lowest *p*-value observed, *p*^*^, is recorded. When comparing the case and the control sets, 0.1 * *p*^*^ is used as a threshold, and only genes with a *p*-value lower than 0.1 * *p*^*^ are considered significant.

SCDE reports a *z*-score which we transform to a *p*-value using the formula *p* = 2Φ(–|*z*|), where Φ(*x*) is the cumulative density of the standard normal distribution. When choosing the threshold for SCDE, we used the same strategy as for D^3^E.

For DESeq2 we used the adjusted *p*-value reported by the algorithm, and we required it to be < .1 to be significant.

### Calculating auto-correlation times

The power spectral density, *S*(*ω*), of the mRNAs for the transcriptional bursting model is given by [9] 

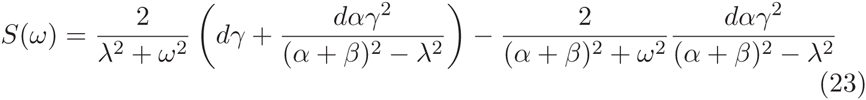
where *d* = *α*/(*α* + *β*). By definition, the auto-correlation, *R*(*t*), is given by the inverse Fourier transform of *S*(*ω*),

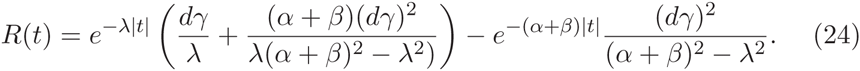

The characteristic time of the auto-correlation is defined as *τ_c_* = *S*(0)/2*R*(0).

## Author’s contributions

MH conceived the study and supervised the research, MD and MH carried out the research, MD wrote the code, and MD and MH jointly wrote the manuscript.

## Acknowledgements

The authors would like to thank Tallulah Andrews, Daniel Gaffney, Vladimir Kiselev, Michael Kosicki, John Marioni, and Gosia Trynka for helpful discussions and comments on the manuscript. MD was funded by the University of Cambridge BBSRC DTP, and MH was funded by the Wellcome Trust.

## Figures

### Additional Files

**Additional file 1 — Figure S1.** Sensitivity to changes in parameters of the Poisson-Beta distribution for the Cramér-von Mises, KS and likelihood ratio tests similar to Fig. 4. The range over which the parameters are varied is larger here. The increased presence of lighter colors shows that for the most part it is easier to identify DE genes for this set of parameters.

**Additional file 2 — Figure S2.** Change in expression levels for 90 genes from the 2-cell and 4-cell mouse embryos as quantified using either qPCR or RNA-seq [5].

**Additional file 3 — Table S1.** Expression levels for the 90 genes from the 2-cell and 4-cell mouse embryos as quantified using either qPCR or RNA-seq [5]. The last six columns indicate the genes that were identified as differentially expressed by different DE algorithms as well as a t-test for the qPCR data.

**Additional file 4.** Parameters for the Islam et al. data without degradation rates. Parameters for the 12,135 genes that were expressed in both cell types.

**Additional file 5.** Parameters for the Islam et al. data with degradation rates. Parameters for the 2,105 genes that were expressed in both cell types, and where degradation rates were available.

**Additional file 6 — Figure S3.** Scatterplots showing the mean fold-change, as well as the fold-change of the CV compared to the change in degradation rate, burst frequency, duty cycle, burst size. In all panels, black dots represent genes which did not change, red dots represent genes which were deemed significant by D^3^E.

**Additional file 7 — Figure S4.** Scatterplots showing the mean fold-change and the fold-change of the characteristic time, as well as the fold-change of the characteristic time compared to the change in degradation rate, variance and characteristic promoter time. In all panels, black dots represent genes which did not change, red dots represent genes which were deemed significant by D^3^E.

**Additional file 8 — Figure S5.** Scatterplots showing the mean fold-change, as well as the fold-change of the mean compared to the change in burst frequency, duty cycle, burst size for early vs late blastocysts from Deng *et al* [10]. In all panels, black dots represent genes which did not change, red dots represent genes which were deemed significant by D^3^E.

**Additional file 9 — Figure S6.** Scatterplots showing the mean fold-change, as well as the fold-change of the CV compared to the change in burst frequency, duty cycle, burst size for early vs late blastocysts from Deng *et al* [10]. In all panels, black dots represent genes which did not change, red dots represent genes which were deemed significant by D^3^E.

